# Regional distribution of white matter hyperintensity burden in coronary artery disease and links with coronary revascularization procedure

**DOI:** 10.64898/2026.05.12.724587

**Authors:** Zacharie Potvin-Jutras, Stéfanie A. Tremblay, Ali Rezaei, Safa Sanami, Dalia Sabra, Brittany Intzandt, Lindsay N. Wright, Christine Gagnon, Amélie Mainville-Berthiaume, Olivier Parent, Mahsa Dadar, Josep Iglesies-Grau, Christopher J. Steele, Mathieu Gayda, Anil Nigam, Louis Bherer, Claudine J. Gauthier

## Abstract

**Introduction:** Coronary artery disease (CAD) increases the risk of cerebrovascular events, yet early brain injury in this population remains poorly characterized. White matter hyperintensities (WMHs), a biomarker of cerebrovascular lesions, are prevalent in CAD and are linked to risk of stroke. Beyond total burden, spatial distribution of WMHs carries pathological significance and is critical for understanding CAD-related injury. While clinical outcomes including coronary revascularization procedure and myocardial infarction influence CAD prognosis, their impact on WMH burden remains unclear.

**Methods:** This study investigated regional WMH burden in CAD and its relationship with clinical characteristics. 82 adults over 50 years participated, including 44 individuals with CAD and 38 controls. WMHs were segmented from fluid attenuated inversion recovery and T1-weighted MRI and categorized as total, periventricular, deep, and superficial regions. History of myocardial infarction and coronary revascularization (coronary artery bypass grafting (CABG) and percutaneous coronary intervention (PCI)), was obtained from medical files.

**Results:** Individuals with CAD exhibited higher total, periventricular, and deep WMH volumes than controls. Participants who underwent CABG had higher superficial WMH volumes than those with PCI, suggesting greater disease severity influences WMH burden.

**Conclusion:** CAD is characterized by a distinct pattern of cerebrovascular vulnerability, with revascularization procedures influencing WMH burden

## Introduction

Coronary artery disease (CAD) is the most prevalent form of heart disease globally (1), affecting approximately 315 million individuals (2). Beyond the elevated risk of acute cardiac events, individuals with CAD also have a higher incidence of ischemic strokes (3) and dementia (4).Importantly, even before the onset of these brain-related conditions, individuals with CAD can show early signs of cerebral vulnerability as measured through brain atrophy (5), perfusion deficits (6,7), and larger white matter (WM) lesion volumes (8).

WM lesions, commonly measured using magnetic resonance imaging (MRI) as white matter hyperintensities (WMHs), are frequently observed in individuals with CAD. (8,9) The etiology of WMHs remains unclear and is likely multifactorial, with evidence implicating chronic cerebral hypoperfusion, blood-brain barrier dysfunction, and cardiovascular risk factors, particularly hypertension. (10) Importantly, WMHs are clinically significant, as they have been linked to cognitive decline, elevated risk of dementia, and stroke. (10,11) Moreover, the anatomical distribution of WMHs is thought to reflect different pathological mechanisms and clinical outcomes, (12,13) with growing evidence that regional WMH burden carries important implications for cognition. (14) Periventricular WMHs are primarily linked to upstream hemodynamic stress, atherosclerosis, elevated blood pressure, and steeper cognitive decline. In contrast, deep WMHs are associated with downstream cerebrovascular damage, small vessel disease, higher body mass index, mood disorders, and, to a lesser extent cognitive difficulties. (13,15–18) However, evidence supporting clearly distinct etiologies between periventricular and deep WMHs remains mixed. Although not commonly assessed, superficial WMHs present significant differences compared to periventricular and deep WMHs, as they indicate more subtle WM tissue alterations and are not associated as consistently with cognitive deficits, although evidence is limited. (12) The regional patterns of WMH burden–spanning periventricular, deep, and superficial WM–and their relationship with CAD remain poorly understood and may help explain the patterns of cognitive deficits associated with CAD.

A history of myocardial infarction (MI) is associated with higher WM lesion load as well as an increased risk of behavioural impairment, cognitive decline, dementia, and stroke. (19,20) Individuals with MI who subsequently undergo coronary artery bypass grafting (CABG) may experience an additional elevation in stroke risk. (21) Moreover, CABG has been linked to the development of new WMHs, likely resulting from microembolization and cerebral hypoperfusion. (22) Thus, despite both MI and coronary revascularization procedures being closely linked to brain health, the extent to which these clinical characteristics influence the regional WMH volumes distribution in individuals with CAD remains poorly understood. To address these gaps, the present study investigates the regional distribution of WMHs in CAD and examines the potential impact of clinical interventions on WMH burden.

## Materials and Methods

### Participants

This cross-sectional study (7,23,24) recruited 99 adults aged 50 years and older. Twelve participants withdrew: two due to claustrophobia during the MRI, five because of interruptions related to the COVID-19 pandemic, two due to discomfort during the MRI, and three for personal reasons (e.g., loss of interest or limited time). Therefore, 87 participants (46 individuals with CAD and 41 healthy controls (HCs)) were included in the data analysis. Prior to statistical analysis, five participants were excluded for the following reasons: one HC due to an MRI-related incidental finding, one HC due to poor FLAIR MRI data quality, one individual with CAD lacked blood pressure information, and two participants (1 individual with CAD and 1 HC) were identified as outliers with total WMH volumes greater than 2.5 standard deviation above the group mean. Thus, statistical analyses were performed on a final sample of 82 participants (44 individuals with CAD and 38 HCs). The study was approved by the Comité d’éthique de la recherche et du développement des nouvelles technologies (CÉRDNT) de l’Institut de Cardiologie de Montréal, in accordance with the Declaration of Helsinki. Written informed consent was obtained at the first visit.

Prior to recruitment, a health history questionnaire was administered on the phone and reviewed during the first visit to determine participants’ eligibility. Medical files were used to collect information on participants’ history of myocardial infarction and type of coronary revascularization procedure, including percutaneous coronary intervention (PCI) and coronary artery bypass grafting (CABG). Inclusion criteria were fluency in either English or French for all participants, a mini-mental state examination (MMSE) and to be 50 years of age or older and absence of neurological or psychiatric conditions. For individuals with CAD, additional inclusion criteria were a confirmed diagnosis of CAD, supported by documentation of acute coronary syndrome, myocardial ischemia, coronary angiography, or a history of coronary revascularization. HCs were only recruited in the absence of cardiovascular conditions, including hypertension and diabetes. Exclusion criteria included a MMSE score below 25 (assessed at the first visit); however, no participants met this criterion. Additional exclusion criteria included a history of stroke, neurological, psychiatric or respiratory disorders, thyroid disease, cognitive impairment, tobacco use or high alcohol consumption (defined as more than 2 drinks per day), contraindications to magnetic resonance imaging (MRI) (e.g., ferromagnetic implant, claustrophobia), current use of oral or patch hormone therapy, exercise limitations affecting cardiorespiratory fitness testing (not used here) and discomfort with hypercapnia (which was added due to the acquisition of data during hypercapnia and not used here). Moreover, participants were excluded if they had a recent acute coronary event (< 3 months), chronic systolic heart failure, resting left ventricular ejection fraction < 40%, symptomatic aortic stenosis, severe nonrevascularizable coronary artery disease (e.g., left main coronary stenosis), awaiting coronary artery bypass surgery, or an implanted automatic defibrillator or permanent pacemaker.

### Magnetic resonance imaging acquisition

Whole-brain MRI data were collected using a 3T Siemens Magnetom Skyra scanner and a 32-channel head coil at the Montreal Heart Institute. T1-weighted images were acquired with a Magnetization Prepared RApid Gradient Echo (MPRAGE) sequence (TR= 2300 ms, TE= 2.32 ms, flip angle= 8°, resolution= 0.9 mm isotropic) to measure total intracranial volume. Axial fluid attenuated inversion recovery (FLAIR) images (TR= 9000 ms, TE= 91 ms, TI= 2500 ms, flip angle= 150°, resolution= 0.9 x 0.9 x 5.0 mm) were acquired to segment WMHs.

### WMH segmentation analysis

WMHs were segmented from the T1-weighted and FLAIR images using the automated and validated Brain tIssue SegmentatiON (BISON) random forest classifier. (25) Preprocessing of the T1-weighted images was conducted prior to the BISON classifier pipeline and included brain extraction via *brain extraction based on nonlocal Segmentation Technique* (BEaST) (26), denoising, intensity nonuniformity correction, and intensity normalization correction, all performed using the MINC toolkit (https://github.com/BIC-MNI/minc-toolkit-v2). Images were then registered to the MNI-ICBM152 template using *advanced normalization tools* (ANTs) (https://github.com/ANTsX/ANTs). More details on the BISON tool usage can be found here (25).

WMH were further segmented into periventricular, deep, and superficial regions. Periventricular WMH were identified in watershed areas by dilating the borders between the anterior, middle and posterior cerebral artery territories from a cerebral arterial territory atlas (27) by 5mm in toward each arterial territory. Superficial WMH were delineated by dilating the cerebral cortex at the grey matter and WM boundaries by 1 mm (28), as segmented by the BISON classifier.Overlapping areas between the periventricular and superficial regions were classified as periventricular WMH. Deep WMH were determined by subtracting the periventricular and superficial masks from the total WM mask. (28) Total intracranial volume was calculated using the *computational anatomy toolbox* (CAT12) within the Statistical Parametric Mapping (SPM) toolbox. (29)

### Statistical analyses

ANCOVA and binomial logistic regressions analyses were conducted to compare the 2 groups for demographic characteristics, cardiovascular measures, and WMH volumes between individuals with CAD and HCs, controlling for age and sex, with total intracranial volume, body mass index (BMI) and mean arterial pressure (MAP) included as additional covariates in the WMH analyses given their established associations with WMH burden (13). Within the CAD group, ANCOVAs were used to examine the difference between clinical characteristics including history of myocardial infarction (yes/no), type of coronary revascularization procedure (PCI vs. CABG) and WMH volumes, controlling for age, sex and total intracranial volume, BMI and MAP. All statistical analyses were performed using WMH volumes normalised with the Yeo-Johnson transformation. To enhance interpretability, the normalised WMH volumes were subsequently standardized using z-scores. To account for multiple comparisons across tests and regions, p-values were adjusted using the Hochberg step-up procedure to control the family-wise error rate. Finally, statistical significance was defined as corrected p-values less than 0.05.

## Results

Participants’ clinical health characteristics of individuals with CAD and HCs are shown in **Table 1**.

**Table 1.**
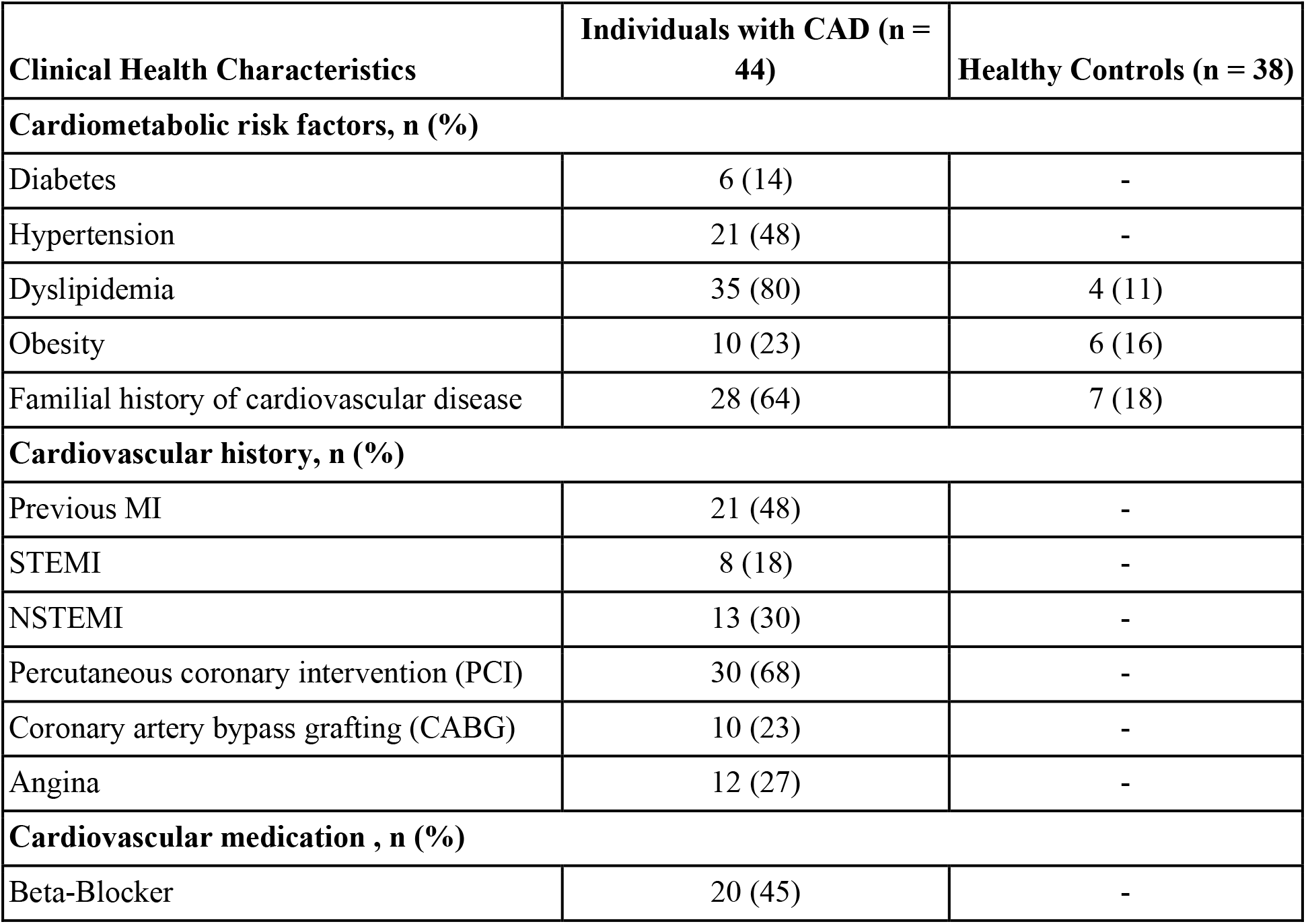
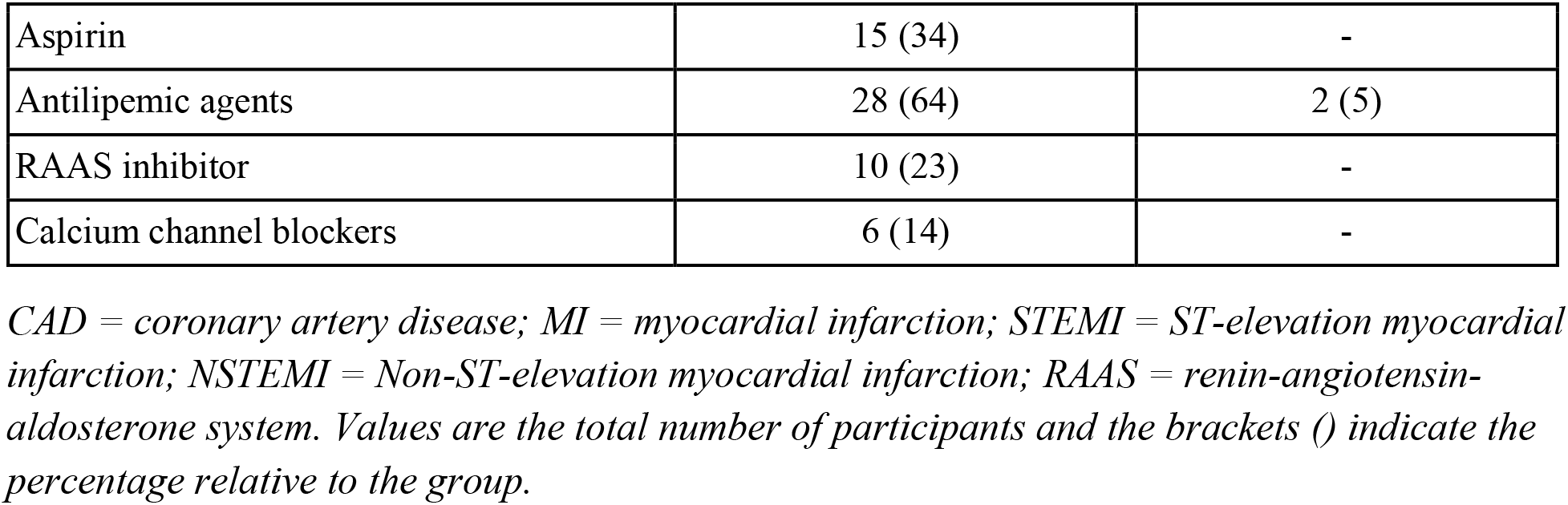
Participants’ characteristics include cardiometabolic risk factors, cardiovascular history and medication of individuals with CAD and HCs.

| Clinical Health Characteristics | Individuals with CAD (n = 44) | Healthy Controls (n = 38) |
| --- | --- | --- |
| **Cardiometabolic risk factors, n (%)** |  |  |
| Diabetes | 6 (14) | - |
| Hypertension | 21 (48) | - |
| Dyslipidemia | 35 (80) | 4 (11) |
| Obesity | 10 (23) | 6 (16) |
| Familial history of cardiovascular disease | 28 (64) | 7 (18) |
| **Cardiovascular history, n (%)** |  |  |
| Previous MI | 21 (48) | - |
| STEMI | 8 (18) | - |
| NSTEMI | 13 (30) | - |
| Percutaneous coronary intervention (PCI) | 30 (68) | - |
| Coronary artery bypass grafting (CABG) | 10 (23) | - |
| Angina | 12 (27) | - |
| **Cardiovascular medication , n (%)** |  |  |
| Beta-Blocker | 20 (45) | - |
| Aspirin | 15 (34) | - |
| Antilipemic agents | 28 (64) | 2 (5) |
| RAAS inhibitor | 10 (23) | - |
| Calcium channel blockers | 6 (14) | - |

*CAD = coronary artery disease; MI = myocardial infarction; STEMI = ST-elevation myocardial infarction; NSTEMI = Non-ST-elevation myocardial infarction; RAAS = renin-angiotensin-aldosterone system. Values are the total number of participants and the brackets () indicate the percentage relative to the group.*

### Group differences between individuals with CAD and HCs

Group differences between individuals with CAD and HCs are displayed in **Table 2**. Individuals with CAD presented significantly higher BMI (p = 0.017; n^2^p = 0.070) compared to HCs.

**Table 2.** Group differences between demographic characteristics and cardiovascular measures.

| **Demographic** | **Individuals with CAD**<br>**(n=44)** | **Healthy Controls**<br>**(n=38)** | **p-values** | **Effect Size ($n^2p$)** |
| --- | --- | --- | --- | --- |
| Age (years) | 67.8 ± 8.8 | 64.9 ± 8.3 | 0.131 | 0.029 |
| Sex (M/F) | 36 / 8 | 28 /10 | 0.352 | 0.011 |
| BMI (kg/m2) * | 28.2 ± 4.4 | 26.2 ± 3.8 | **0.017** | 0.070 |
| Education (years) | 16.0 ± 4.6 | 16.0 ± 2.7 | 0.913 | < 0.001 |
| MMSE score | 28.4 ± 1.2 | 28.2 ± 1.5 | 0.227 | 0.020 |
| **Cardiovascular Measures** |  |  |  |  |
| Systolic BP (mmHg) | 126.3 ± 14.0 | 127.9 ± 14.1 | 0.647 | 0.003 |
| Diastolic BP (mmHg) | 73.4 ± 7.2 | 76.5 ± 9.1 | 0.659 | 0.003 |
| MAP (mmHg) | 91.1 ± 7.6 | 93.6 ± 9.7 | 0.128 | 0.029 |
| Resting Heart Rate (bpm) | 63.5 ± 10.6 | 67.9 ± 10.7 | 0.130 | 0.029 |

*CAD: coronary artery disease; BMI: body mass index; MMSE: mini-mental state examination; BP: blood pressure; MAP: mean arterial pressure. Values are mean ± SD. P-values indicate ANCOVA analyses between individuals with CAD and healthy controls adjusting for age and sex. An asterisk (\*) indicates a significant group difference after Hochberg correction.*

Individuals with CAD showed significantly higher total (p = 0.005; n^2^p = 0.023), periventricular (p = 0.001; n^2^p = 0.053) and deep (p = 0.010; n^2^p = 0.012) WMH volumes compared to HCs (**Figure 1**). For reference, the average WMH volume values prior to normalisation can be found by group in **Table S1** and by sex in **Table S2**.

**Figure 1.**
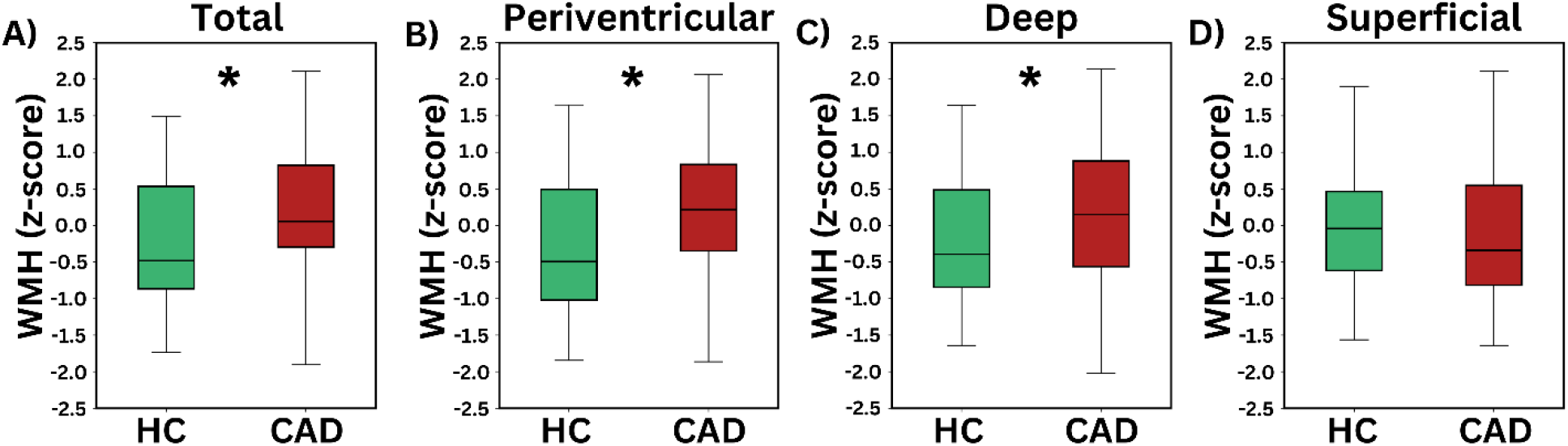
Regional WMH group differences between individuals with CAD and HCs. Box plots from ANCOVA analyses demonstrating group differences in standardized WMH (z-scores) between individuals with CAD (n = 44) and healthy controls (n = 38) in A) total WMH, B) periventricular WMH, C) deep WMH and D) superficial WMH. An asterisk (*) indicates a significant group difference after Hochberg correction.

### Clinical characteristics and WMH volumes in CAD

Among individuals with CAD, surgery type was significantly associated with WMH volume, with the CABG group showing higher superficial WMH compared to the PCI group (p = 0.037; n^2^p = 0.129) (**Figure 2**). No significant effect of having had a MI history on WMH volumes was observed.

**Figure 2.**
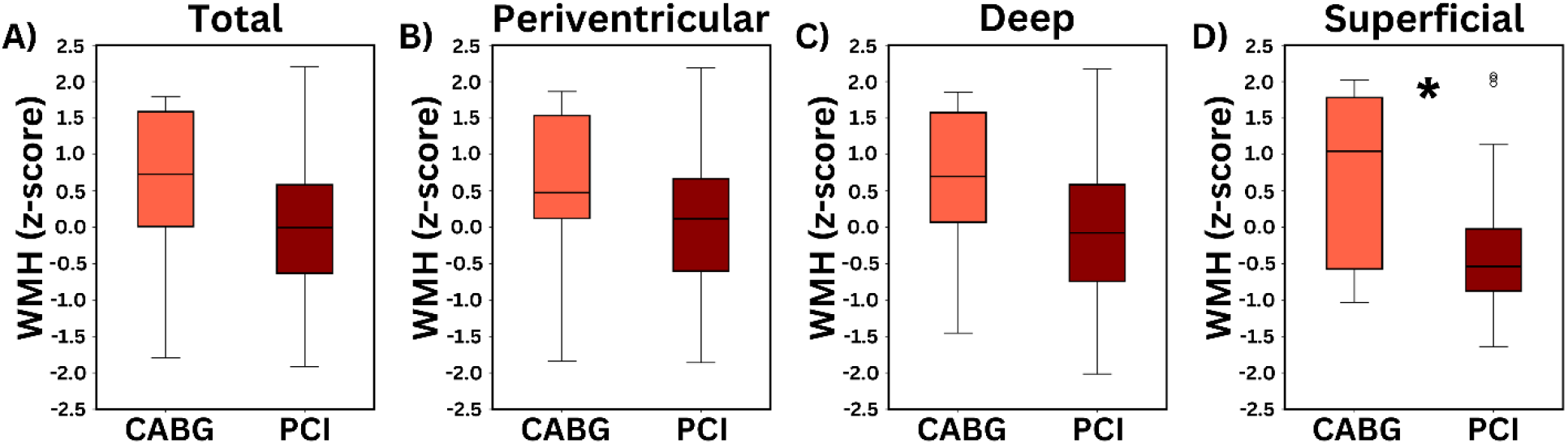
Regional WMH by coronary revascularization procedure type in individuals with CAD. Box plots from ANCOVA analyses showing subgroup differences by coronary revascularization procedure type (CABG (n = 10) vs. PCI (n = 30)) on standardized WMH across A) total WMH, B) periventricular WMH, C) deep WMH and D) superficial WMH. CABG group has significantly higher superficial WMH compared with PCI group. An asterisk (*) indicates a significant group difference after Hochberg correction.

## Discussion

In this study, we investigated WMH volumes in individuals with CAD and their relationship with clinical disease characteristics. Compared to HCs, individuals with CAD exhibited significantly higher BMI, alongside greater total, periventricular, and deep WMH volumes. The predominance of periventricular and deep WMHs suggests region-specific vulnerability in CAD. Although a history of MI was not associated with WMH burden, the type of coronary revascularization procedure influenced WMH distribution, with individuals who had undergone CABG demonstrating greater superficial WMH volumes than those treated with PCI. Taken together, these findings indicate that CAD is linked to regionally-specific patterns of WMH accumulation, highlighting the importance of considering WMH location when interpreting cerebrovascular injury and associated risk.

### WMH burden in CAD

Our findings revealed that individuals with CAD demonstrate greater total, periventricular, and deep WMH volumes compared to HCs. While prior studies have reported elevated overall WMH burden in those at risk for (30,31) or living with CAD (8,9), lesion location has been little examined. To our knowledge, this is the first study to show that periventricular and deep–but not superficial–WMHs are specifically higher in CAD. Notably, group differences were more pronounced for periventricular WMHs compared to deep WMHs. This regional predominance aligns with prior work indicating that periventricular WMHs are particularly sensitive to cardiovascular pathology (31–33), including atherosclerosis (15). In individuals at risk of CAD, periventricular WMHs show stronger association than deep WMHs with coronary artery plaque accumulation (32,33) and coronary artery calcification (31). Furthermore, in a large population study of older adults, several CAD-related genetic variants were more strongly linked to periventricular than deep WMHs. (34) Importantly, periventricular WMHs have also shown greater sensitivity to cognitive decline (35), all-cause dementia (36) and ischemic stroke (37), suggesting that these lesions may serve as an early and location-specific marker of cerebral vulnerability in CAD. Nonetheless, the significantly higher proportion of deep WMHs observed in our CAD cohort proposes multifactorial WM injury, as periventricular and deep WMHs are thought to arise, at least in part, from potentially distinct pathophysiological mechanisms. (13)

### Periventricular WMHs in CAD

In this study, periventricular WMHs were located within watershed areas at the junction of the anterior, middle, and posterior cerebral artery territories. Watershed regions in WM are particularly susceptible to ischemic injury because they are supplied by the most distal arterial branches and are therefore highly vulnerable to hypoperfusion. (38) The presence of periventricular WMHs is thus consistent with the reduced cerebral blood flow previously reported in individuals with CAD by our group and others (6,7), including within this cohort. Pathologically, periventricular WMHs reflect a combination of structural alterations, including axonal degeneration, demyelination, astrocyte proliferation, and ependymal lining damage. (39) Notably, our prior work demonstrated that WM microstructural abnormalities in this CAD cohort were primarily driven by myelin-sensitive imaging techniques. (23) Collectively, these findings suggest that chronic hypoperfusion (7) and myelin loss (23) may represent relevant biological targets for mitigating periventricular WMH accumulation in CAD, although longitudinal mechanistic studies are needed to confirm causal relationships.

### Deep WMHs in CAD

Deep WMHs were also significantly elevated in individuals with CAD, although group differences were less pronounced than those observed for periventricular lesions. The present study defined deep WMHs as lesions located between periventricular watershed territories and superficial WM regions. Deep WMHs may be particularly vulnerable to hypoperfusion, similar to watershed territories, although this may arise from their relatively sparse vascular supply and fewer penetrating arterial branches. (40) Consequently, the reduced CBF previously observed in this CAD cohort may contribute to injury extending beyond periventricular watershed regions into deeper WM structures. Emerging pathological evidence characterizes deep WMHs as lesions showing more severe axonal degeneration, vacuolation, gliosis and demyelination. (39) The presence of demyelination within both deep and periventricular WMHs further supports the role of myelin integrity as an important potential biomarker of WM health in CAD, consistent with findings previously identified within this CAD cohort. (23) Nonetheless, the precise pathological distinctions between periventricular and deep WMHs remains unclear and may vary across disease contexts. In CAD, our results suggest that WM injury involves both regions, highlighting the need for future studies to clarify the shared and distinct processes underlying deep WMHs development.

### Coronary Revascularization Procedures

Within the CAD group, individuals who underwent CABG had higher superficial WMH volumes than those who underwent PCI. CABG is a more invasive procedure than PCI, commonly involving cardiopulmonary bypass and extensive aortic manipulation, both of which increase the risk of perioperative complications such as stroke. (41,42) Postoperatively, CABG has been associated with the development of new WM lesions. (22,43–45) These lesions are often small and therefore difficult to detect, and are hypothesized to stem from a combination of cerebral hypoperfusion and microembolic events occurring during the surgery. (22,43,45,46) The anatomical distribution of these postoperative WMHs remain incompletely characterized. Previous studies describe a relatively spare lesion pattern that frequently includes WM regions adjacent to the cortex (22,43,45), consistent with our observation of greater lesion burden within superficial WM. Superficial WMHs are thought to reflect less severe tissue injury than periventricular or deep WMHs, potentially because they are more strongly influenced by increased water content and subtle microstructural alterations that may be partially reversible. (12) Although superficial WMHs themselves have not been consistently linked to cognitive dysfunction (12), CABG has independently been associated with an increased risk of postoperative cognitive impairment (47). Overall, our findings suggest that individuals with CAD who undergo CABG may be particularly susceptible to the accumulation of superficial WMHs. However, the preferential localization of these lesions may indicate milder tissue alterations, potentially reflecting distinct underlying mechanisms and clinical significance relative to periventricular and deep WMHs.

### Limitations

This study has several limitations. First, our sample included a higher proportion of males compared to females, which limited our ability to investigate sex-specific effects. However, this imbalance partly reflects the lower prevalence of CAD in females, especially among adults younger than 65 years of age. (48) Nonetheless, future studies are necessary to further investigate the effects of sex on regional WMH burden in CAD. Second, many participants with CAD had previously completed cardiac rehabilitation, indicating that our cohort primarily represents clinically stable individuals with CAD. As such, the findings may not fully generalize to individuals with more acute or severe CAD disease presentations. Third, some HCs may have underlying, untreated vascular risk factors, as not all HCs were under regular medical supervision. Finally, the cross-sectional design precludes causal inferences regarding the relationship between clinical characteristics and WMH volumes.

## Conclusions

In summary, individuals with CAD exhibited higher total, periventricular, and deep WMH volumes compared to HCs, suggesting heightened cerebral vulnerability. Moreover, participants with CAD who underwent CABG showed greater superficial WMH volumes relative to individuals treated with PCI. Together, these findings indicate that CAD is associated with distinct spatial patterns of WMH burden that may carry important implications for long-term cerebrovascular and cognitive health. Future mechanistic longitudinal studies and clinical trials are needed to determine whether clinical interventions can mitigate WMH accumulation in CAD and to clarify how specific clinical and cardiovascular factors influence WM integrity over time.

## Acknowledgements

We would like to acknowledge the contributions of Paule Samson, Hakima Benhalima, Julie Lalongé, and all members involved with the project from the ÉPIC Center. We would like to acknowledge research assistants and students who took part in the data acquisition including Roni Zaks, Robert Hovey, Stephanie Beram, Alexandre Bailey, Agathe Godet, Milla Shakleva, Victoria D’Amours, Kathia Saillant, Catherina Medeiros, and Zineb Rouabah.

## Authors Contribution

Z.P.J.: performed experiments, analyzed data, interpreted results of experiments, prepared figures, drafted manuscript, edited and revised manuscript, approved final version of manuscript; S.A.T.: performed experiments, edited and revised manuscript, approved final version of manuscript; A.R.: performed experiments, edited and revised manuscript, approved final version of manuscript; S.S.: performed experiments, edited and revised manuscript, approved final version of manuscript; D.S.: performed experiments, edited and revised manuscript, approved final version of manuscript; B.I.: performed experiments, edited and revised manuscript, approved final version of manuscript; L.N.W.: performed experiments, edited and revised manuscript, approved final version of manuscript; C.G.: performed experiments, edited and revised manuscript, approved final version of manuscript; A.M.B.: performed experiments, edited and revised manuscript, approved final version of manuscript; O.P.: edited and revised manuscript, approved final version of manuscript; M.D.: edited and revised manuscript, approved final version of manuscript; J.I.G: performed experiments, edited and revised manuscript, approved final version of manuscript; C.J.S.: edited and revised manuscript, approved final version of manuscript; M.G.: performed experiments, edited and revised manuscript, approved final version of manuscript; A.N.: performed experiments, edited and revised manuscript, approved final version of manuscript; L.B. conceived and designed research, edited and revised manuscript, approved final version of manuscript; C.J.G.: conceived and designed research, performed experiments, interpreted results of experiments edited and revised manuscript, approved final version of manuscript.

## Ethics approval and informed consent statements

The study was approved by the Comité d’éthique de la recherche et du développement des nouvelles technologies (CÉRDNT) de l’Institut de Cardiologie de Montréal, in accordance with the Declaration of Helsinki. Written informed consent was obtained at the first visit.

## Data availability statement

Anonymized and defaced data that support the findings of this study will be made openly available on OpenNeuro (https://openneuro.org) within 6 months of publication. A persistent DOI will be provided upon release.

## Supplementary Materials

Table S1 and Table S2 can be found in the supplementary materials.

