## Supplemental Table 1 & 2 for "Regional distribution of white matter hyperintensity burden in coronary artery disease and links with coronary revascularization procedure"

**Supplementary Materials**

**Table S1. Group differences in WMH volumes prior to normalization**

|  | **Individuals with CAD (n=44)** | **Healthy Controls (n=38)** |
| --- | --- | --- |
| **Total Intracranial Volume (cm3)** | 1527.09 ± 118.29 | 1566.87 ± 176.41 |
| **White Matter Hyperintensities Volumes (cm3)** | | |
| Total | 6.72 ± 9.79 | 3.94 ± 1.91 |
| Periventricular | 3.41 ± 5.07 | 2.08 ± 2.52 |
| Deep | 2.50 ± 4.04 | 1.48 ± 1.53 |
| Superficial | 0.82 ± 1.84 | 0.39 ± 0.37 |

*CAD = coronary artery disease*

**Table S2. Sex differences in WMH volumes prior to normalization**

|  | **Individuals with CAD (n = 44)** | | **Healthy Controls (n = 38)** | |
| --- | --- | --- | --- | --- |
|  | Females (n = 8) | Males (n = 36) | Females (n = 10) | Males (n = 28) |
| **Total Intracranial Volume (cm³)** | 1413.88 ± 81.01 | 1552.25 ± 110.89 | 1383.10 ± 144.71 | 1632.50 ± 136.71 |
| **White Matter Hyperintensities Volumes (cm³)** | | | |  |
| Total | 14.10 ± 19.03 | 5.09 ± 5.50 | 2.39 ± 2.48 | 4.50 ± 4.54 |
| Periventricular | 7.72 ± 9.82 | 2.46 ± 2.66 | 0.94 ± 0.85 | 2.48 ± 2.79 |
| Deep | 5.48 ± 8.02 | 1.83 ± 2.18 | 1.08 ± 1.13 | 1.62 ± 1.65 |
| Superficial | 0.90 ± 1.32 | 0.80 ± 1.95 | 0.37 ± 0.56 | 0.39 ± 0.29 |

*CAD = coronary artery disease*
